# Co-infection of Ehrlichia with *B. burgdorferi* drives emergency myelopoiesis and promotes Lyme arthritis

**DOI:** 10.1101/2022.08.23.505055

**Authors:** Jesse L. Bonin, Steven R. Torres, Ashley L. Marcinkiewicz, Xiuli Yang, Utpal Pal, Julia M. DiSpirito, Tristan Nowak, Yi-Pin Lin, Katherine C. MacNamara

## Abstract

Lyme disease is caused by the extracellular pathogen *Borrelia burgdorferi* (*Bb*), transmitted by the *Ixodes scapularis* tick. Approximately one-third of infected individuals develop arthritis of weight-bearing joints, though it is unclear why some patients develop arthritis and severe systemic disease while others do not. C57BL/6 (B6) mice are susceptible to *Bb* infection but do not develop arthritis, providing an *in vivo* model to evaluate mechanisms regulating development of Lyme arthritis. We demonstrate here that co-infection of B6 mice with the tick-borne pathogens *Bb* and *Ehrlichia muris* (*Em*) induced significant arthritis. Although co-infection did not impact bacterial burden or growth of either pathogen, the resultant Lyme arthritis in co-infected mice correlated with significant hematologic disturbances. Whereas single *Bb* infection elicited no overt hematologic changes, *Em* infection resulted in thrombocytopenia, lymphopenia, monocytosis, and granulocytosis, which was consistently observed in mice co-infected with both *Bb* and *Em*. Hematologic changes correlated with profound changes to the hematopoietic stem and progenitor cell (HSPC) populations in *Em*-infected mice. Most notable were dramatic reductions in populations of HSPCs committed to myeloid-biased differentiation. Co-infection resulted in persistent hematologic changes and bone marrow inflammation. Our data demonstrate for the first time that B6 mice, resistant to developing Lyme arthritis, exhibit severe joint pathology in the presence of a second pathogen, correlating with persistent emergency myelopoiesis. Our data support the conclusion that pathogen burden is not sufficient for disease and specific inflammatory signals and cells regulate the development of Lyme arthritis.

**Importance:** Tick-borne illnesses, historically relegated to specific geographic areas, are increasing in prevalence and distribution. *Borrelia burgdorferi* causes Lyme disease, the most common tick-borne illness in North America, characterized by debilitating arthritis, carditis, and neurologic complications. It remains unclear why certain infected individuals develop severe disease while others are only mildly symptomatic. Human monocytic ehrlichiosis (HME) is another tick-borne disease that often results in profound illness and severe hematological disturbances. We show here that co-infection of B6 mice, resistant to Lyme arthritis, with *Borrelia burgdorferi* and *Ehrlichia muris*, used to model HME, results in the development of severe arthritis and emergency myelopoiesis. Our work suggests that immune activation driven by co-infection contributes to the development of Lyme arthritis.

## Introduction

*Borrelia burgdorferi* sensu stricto (*Bb*) is the main cause of Lyme disease in North America (1, 2). Originally detected in Lyme, Connecticut, incidence of Lyme borreliosis has increased dramatically in prevalence and geographic distribution over the last four decades (3), with an estimated 476,000 cases per year in North America (2). After transmission to humans via *Ixodes* ticks (1, 4), infection with *Bb* often results in a characteristic erythema migrans rash, the result of a localized immune response to the bacterium. Dissemination of the bacterium from the site of infection to distal tissues including joints results in arthritis of the weight-bearing joints in 27.5 % of infections (5); it remains unclear why only some patients develop arthritis. The bacterium itself does not produce toxins (6), and recruitment of immune cells, notably neutrophils, is thought to cause significant inflammation of the joint, contributing to arthritis (7, 8). The immune response to *Bb* is driven by recognition of bacterial lipoproteins by host TLRs. *In vitro* infection of human PBMCs results in production of TNF, IL-6, IL-10, and IL-1β (9). Systemic effects of *Bb*, as exhibited by changes to circulating immune cells, are not observed in patients with Lyme arthritis, which may be a result of the pathogen’s ability to suppress or evade acute inflammatory responses (10).

In addition to Lyme disease-causing bacteria, *Ixodes* ticks carry a variety of pathogens that infect humans. Human monocytic ehrlichiosis (HME) results from infection with the tick-borne intracellular pathogen *Ehrlichia chaffeensis*, spread by the *Amblyomma americanum* tick (11), and *E. muris* eauclarensis, carried by *I. scapularis* (12). Important clinical signs of HME include hematologic disturbances, characterized by significant reductions in circulating platelets (thrombocytopenia) and lymphocytes (lymphopenia) (13). *Ehrlichia muris* (*Em*) is closely related to *E. chaffeensis* (14) and causes disease in mice similar to what is observed in human patients with HME. Co-infection of mice with *Bb* and *Babesia microti* resulted in alterations to the immune response against both pathogens, with more severe Lyme arthritis and lower *Babesia microti* burdens in co-infected mice (15, 16). A study comparing the signs and symptoms of patients with Lyme disease with those of patients infected with other tick-borne pathogens revealed that hematologic disturbances were absent in Lyme disease, as compared to other tick-borne infections including babesiosis, anaplasmosis, and ehrlichiosis, where profound hematological changes were noted (17). Furthermore, when hematological abnormalities were observed in patients with borreliosis, it was found to correlate with the presence of a co-infecting pathogen. Concurrent Lyme borreliosis with babesiosis or ehrlichiosis resulted in a wider variety of symptoms in infected patients, and more flu-like symptoms than those with Lyme borreliosis alone (17). Moreover, in 39% of suspected tick-borne infections, multiple pathogens were present (17). Therefore, co-infection poses a challenge to both identification and treatment of tick-borne diseases.

Human monocytic ehrlichiosis (HME) induces profound changes to the hematologic parameters of infected individuals (18). *Em* is used in mice to model HME, and causes disruption of the hematopoietic stem cell niche **(19–21)**. Hematopoietic stem and progenitor cells (HSPCs) give rise to all differentiated blood cells in adult mammals through the process of hematopoiesis. HSPCs typically reside in the bone marrow and are maintained by dynamic signaling processes within the marrow microenvironment (22, 23). Under homeostatic conditions, hematopoiesis is regulated through endogenous signaling from stromal and hematopoietic cells, however blood production can be altered during infection or inflammation, which can promote the generation of immune cells necessary for dealing with microbial challenge **(24–26)**. Thus, how co-infecting tick-borne pathogens impact hematopoiesis needs to be carefully studied to further to our understanding of pathogenesis and transmission of vector-borne infections.

To address the impact of bacterial co-infection on pathogenesis of *Bb*-induced disease, we used C57BL/6 (B6) mice to study co-infection of *Bb* with *Em*. Consistent with previous observations, B6 mice were resistant to developing *Bb*-induced Lyme arthritis and did not experience a significant immune response to the pathogen (27, 28). Co-infection of *Bb* with *Em*, however, resulted in the development of marked arthritis of the ankle joint. These changes did not correlate with changes in the persistence of *Bb* or *Em*. However, hematologic parameters and inflammatory cytokine production of co-infected mice resembled that of *Em*-infected mice. Our data reveal robust demand-adapted myelopoiesis as evidenced by significantly increased circulating monocytes and granulocytes, reduced circulating lymphocytes, and profound thrombocytopenia. Moreover, within the bone marrow we observed a profound loss of HSPCs in the context of *Em* infection, whereas *Bb* was not associated with changes to the HSPC compartment. Taken together, our work identifies a role for co-infection in driving changes to blood cell production that are linked with the development of Lyme arthritis. These studies demonstrate that hematopoietic programs may underlie the progressive and periodic nature of Lyme arthritis.

## Methods

### Mice

Female, 6–8-week-old C57BL/6 (B6NTac) mice were purchased from Taconic (Germantown, NY). All animal experiments were performed following approval by the IACUCs at AMC (ACUP 20-04004) and Wadsworth (Protocol docket numbers 16-431, 19-451, 22-451).

### Infection of mice and xenodiagnosis

Mice were infected via i.p. injection of bacteria in sucrose-phosphate-glutamate (SPG) buffer. Mice were inoculated with 100,000 copies each of *Em* and *Bb. Em* stocks were generated from infected splenocytes as previously described (29). *Bb* strain B31-A3 was grown in Barbour-Stonner-Kelly (BSK)-II compete medium at 33° C (30). Prior to infection, cultured *B. burgdorferi* was analyzed with PCR to confirm the presence of all plasmids (31). Mice were sacrificed at 10 or 22 days post infection (dpi) and tissues collected.

Mice used for xenodiagnosis were i.p. infected as described above. At 22 dpi, approximately 100-150 naïve, unfed *Ixodes scapularis* larvae (cultivated from engorged, field-collected adults that were assessed to be specific pathogen free) were placed on the mice and allowed to feed until repletion. Ticks were collected and frozen at -20°C until processed for analysis.

### Bacterial burden

Mouse tissues for bacterial burdens were collected at both 10 and 22 dpi. DNA was extracted from murine spleens using an EZNA Tissue DNA kit (Omega Bio-Tek, Georgia, USA). *E. muris* burden in infected mice was quantified using spleens via RT-qPCR using a Mastercycler ep *Realplex* (Eppendorf, Hamburg, Germany) (32) (Table S1). To test for *Bb* burdens, DNA from mouse ears and bladder were extracted using an EZ-10 Genomic DNA kit (Biobasic, Markham, ON, Canada). qPCR was then performed to quantify *Bb* loads. Spirochete genomic equivalents were calculated using an ABI 7500 Real-Time PCR System (ThermoFisher Scientific) in conjunction with PowerUp SYBR Green Master Mix (ThermoFisher Scientific) based on amplification of the Lyme borreliae *recA* gene (Table S1) with the amplification cycle as described previously (33). The number of *recA* copies was calculated by establishing a threshold cycle standard curve of a known number of recA gene extracted from cultured B31-A3.

To determine *Bb* burdens in xenodiagnostic ticks, frozen ticks were crushed, and the DNA extracted using an EZ-10 Genomic DNA kit (Biobasic). qPCR was performed on ten ticks per mouse, at a minimum of two technical replicates. Any tick with an average *recA* burden above the lowest standard was considered positive, and any mouse with a minimum of one positive tick was considered to be infected.

### Blood collection for CBC and serum analysis

Blood was collected from mice via cardiac puncture into BD Microtainer Tubes with K_2_EDTA (Cat. No. 365974). CBC analysis was performed using a Heska ElementHT5 Veterinary Hematology Analyzer (Heska Corporation, Loveland, Colorado). Blood for serum analysis was collected via cardiac puncture and spun to separate cells. The levels of total anti-*Bb* IgG in the sera were quantified as previously described (34).

### Protein analysis

Protein from bone marrow was collected my homogenization of bone marrow with a pestle in lysis buffer made with IGEPAL CA-630 and proteinase inhibitors. Analysis of chemokines present in the serum and bone marrow of mice was performed using a Bio-Plex Pro Mouse Chemokine Panel 31-Plex (Bio-Rad, Hercules, California), and for bone marrow, data were normalized to total protein.

### Flow Cytometry

BM cells were isolated from murine tibia and femora via flushing with HBSS, followed by passage through a 70 μm sterile filter, and spleens were processed through crushing between frosted glass slides and passage through a 70 μm filter. Single-cell suspensions underwent red blood cell lysis via ACK buffer, followed by counting and plating. Cells were stained with fluorescently labeled antibodies described in Table S2. Data were collected on a FACS LSR II or FACS Symphony (BD Biosciences, San Jose, California). Data were analyzed using FlowJo v10 (BD, San Jose, California).

### Histopathology

Talocrural (ankle) joints were collected from mice at 22 dpi. Joints were fixed for 48 hours with 10 % neutral-buffered formalin followed by decalcification for one week with 10 % formic acid. These were embedded in paraffin wax and sliced to make slides that were strained with hematoxylin and eosin (Wadsworth Histopathology Core Facility, NYS Department of Health, Albany, NY, USA). Arthritis severity was scored as previously described (34, 35). Briefly, two sections per mouse joint were blindly evaluated for the severity of *Bb*-induced arthritis based on the inflammatory scores of 0 (no inflammation), 1 (mild inflammation with less than two small foci of infiltration), 2 (moderate inflammation with two or more foci of infiltration), or 3 (severe inflammation with focal and diffuse infiltration covering a large area). The infiltration of different cell types (i.e., neutrophils and lymphocytes) were highlighted in high-resolution figures.

### Statistical Analysis

Statistical tests for xenodiagnostic ticks were performed using two-way Fisher’s exact test. Statistical tests between multiple groups were performed using one-way ANOVA with Tukey’s multiple comparisons. All data were analyzed using GraphPad Prism 9 (San Diego, California).

## Results

### *Em* and *Bb* co-infection results in severe Lyme arthritis in B6 mice

C57BL/6 mice can be colonized by *Bb* at high doses but do not develop Lyme borreliosis-associated disease manifestations (e.g. arthritis) (36, 37). To determine the impact of co-infection with other tick-borne pathogens on the overall infection and severity of such manifestations, C57BL/6 mice were i.p. co-infected simultaneously with *Bb* and the tick-borne pathogen *Ehrlichia muris* (*Em*), or singly infected with each of these pathogens (**Fig. 1A**). Mice were euthanized at 10 and 22 days post-infection (dpi), the respective peaks of severity for *Em* and *Bb* infection (8, 19).

**Figure 1.**
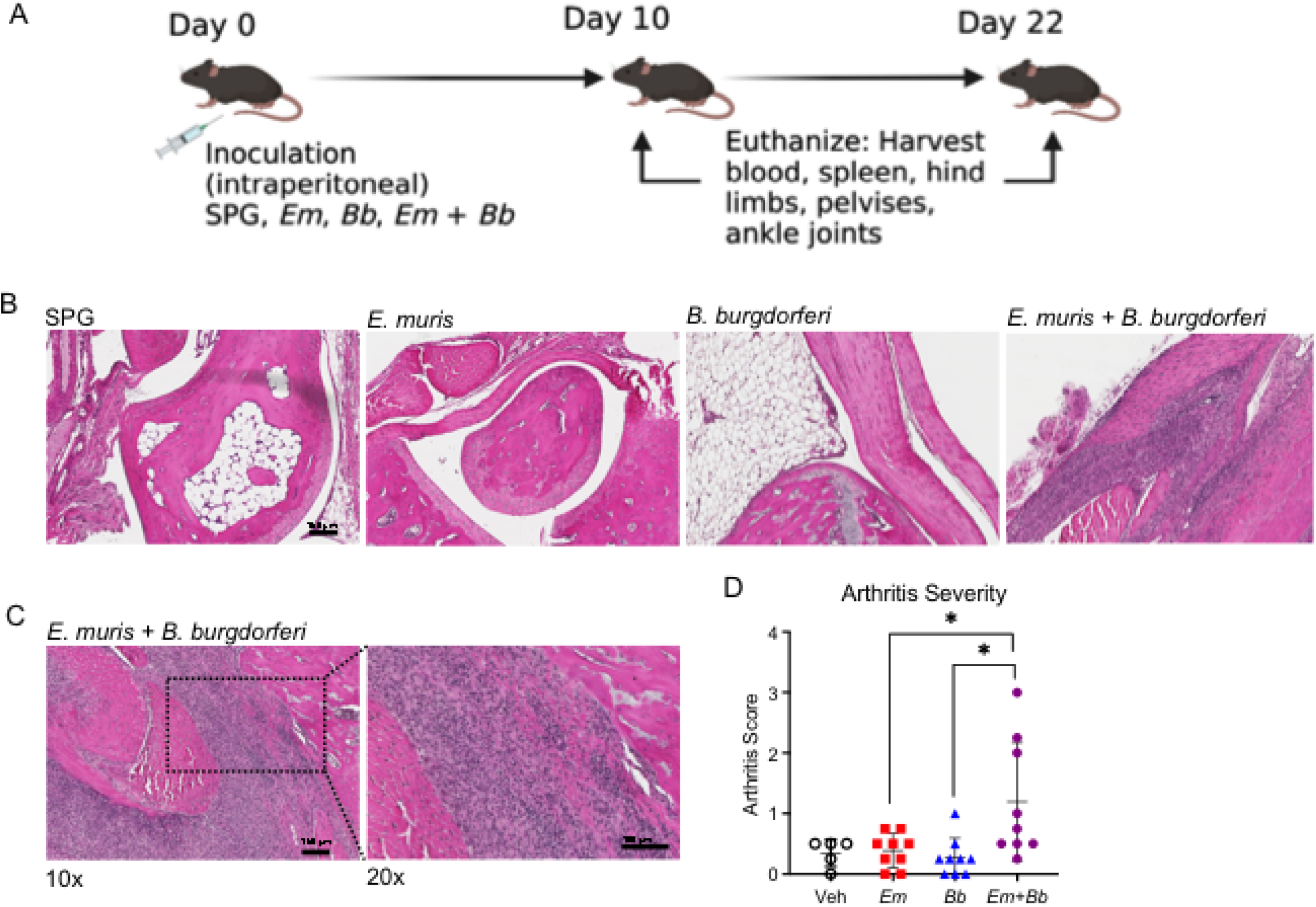
Co-infection of *E. muris* with *B. burgdorferi* induces arthritis. (**A**) Schematic of experiment where Lyme-resistant C57BL/6Tac mice were infected with the tick-borne pathogens *E. muris* and *B. burgdorferi*. Mice were euthanized at 10- or 22-days post infection for analysis of tissues. Panel created with Biorender.com. (**B**) Histological analysis of talocrural joints was performed at 22 dpi on specimens stained with hematoxylin and eosin for vehicle control (SPG) mice and mice infected with *Em, Bb*, or both *Em* and *Bb*. (**C**) Additional images including higher magnification image from *Em* and *Bb* co-infected mice are shown. (**D**) Samples were scored to determine arthritis severity. Data points represent individual mice; error bars represent standard deviation. Groups were compared using a one-way ANOVA with Tukey’s post-hoc comparison. *p<0.05.

To confirm infection of both pathogens, particularly in the context of co-infected animals, spleens were obtained for determining ehrlichial burdens (19). *Em* was detected in single *Em*-infected and co-infected mice at 10 dpi with similar burdens among the two groups (**Supp. Fig. 1A)**. *Em* was undetectable by 22 dpi in both single- and co-infected groups (data not shown). We then attempted to determine *Bb* burden in multiple tissues (i.e., ears and bladder) at 10 and 22 dpi (early and peak timepoints for borreliosis, respectively (8)) but failed to detect any burdens (data not shown). This is consistent with the fact that mice are less susceptible to *Bb* infection using i.p. infection relative to other routes of infection (i.e., i.d. or tick feeding) (38), and C57BL/6 mice are less susceptible to *Bb* infection (39). We therefore examined sera at 22 dpi to test for *Bb*-specific antibody responses. As expected, anti-*Bb* IgG antibodies were not detectable at 10 dpi (data not shown) but were present in single-*Bb* and co-infected mice at 22 dpi, with no difference in the titers between these groups (**Supp. Fig. 1B**).

We next determined the severity of arthritis in infected mice. Hematoxylin and eosin staining of the talocrural joint at 10 dpi, the peak of *Em* infection (19) revealed little to no inflammation in the joints (data not shown). At 22 dpi, the respective peak for *Bb* infection (8), there were minimal changes to the joints of mice infected with *Em* or *Bb* alone, whereas the co-infected mice developed severe inflammation around the tendons and synovial membrane (**Fig. 1B-C**). This infiltration was consistent with significantly higher arthritis scores of the joints from co-infected mice compared to mock and single-infected mice (**Fig.1D**).

To confirm that mice were indeed infected with *Bb*, we performed a xenodiagnostic assay, a rigorous and sensitive strategy to identify infection with *Bb* (40– 42). Naïve, flat larval ticks were placed on infected mice at 22 dpi (8) and after feeding to repletion were analyzed to test for the presence of *Bb* genetic material. Whereas the mock and singly-*Em* infected mice yielded no positive ticks, all singly-*Bb* and co-infected mice resulted in at least one tick per mouse was positive for *Bb*, indicating that all mice in the single and co-infected groups were indeed infected and colonized by viable *Bb* (**Supp. Fig. 1C**). Although not a direct measure of bacterial burden in mice, these data together with antibody titers demonstrate that B6 mice were able to support *Bb* infection. Furthermore, the presence of *Em* in co-infected mice was not necessary for *Bb* infection. Together, our data demonstrate the *Bb-*infection of Lyme-arthritis-resistant B6 mice develop inflammation of the joints when co-infected with *Em*.

### *Em* co-infection drives a myeloid-biased immune response

The hematologic parameters of *Bb*-infected individuals are often unremarkable (17, 43), while ehrlichiosis results in significant hematologic disturbances (21). We therefore investigated whether the hematologic parameters of co-infected mice differed significantly from singly-infected mice at the peak of *Em* infection (10 dpi (19)). Complete blood counts (CBC) revealed that the co-infected mice resembled single *Em*-infection, with significantly increased myeloid cells and profound lymphopenia (**Fig. 2A**). Co-infected mice also exhibited significant thrombocytopenia, consistent with what was seen in *Em* infection (**Fig. 2B**), although only mild anemia was seen in all infected mice (**Fig. 2C**). Mice infected with *Em* exhibited profound lymphopenia, and this was not impacted by co-infection with *Bb* (**Fig. 2D**), although co-infection resulted in slightly more significant thromobocytopenia than single *Em* infection. Granulocytes and monocytes were elevated in both *Em*- and co-infected mice at 10 dpi, while *Bb* infection caused no perturbation in myeloid cells, and closely resembled vehicle-inoculated controls (**Fig. 2E-H**). These data suggest that the immune response in co-infected mice is primarily driven by responses to *Em*. Mice co-infected with both *Em* and *Bb* exhibited significantly elevated neutrophils, monocytes, and basophils relative to singly-*Em*-infected mice, thus demonstrating that co-infection elicited an exacerbated inflammatory response. Together, our findings reveal that the presence of *Em* dominated the early innate immune response and suggest that a lack of inflammatory responses against *Bb* can be overcome by a co-infecting pathogen.

**Figure 2.**
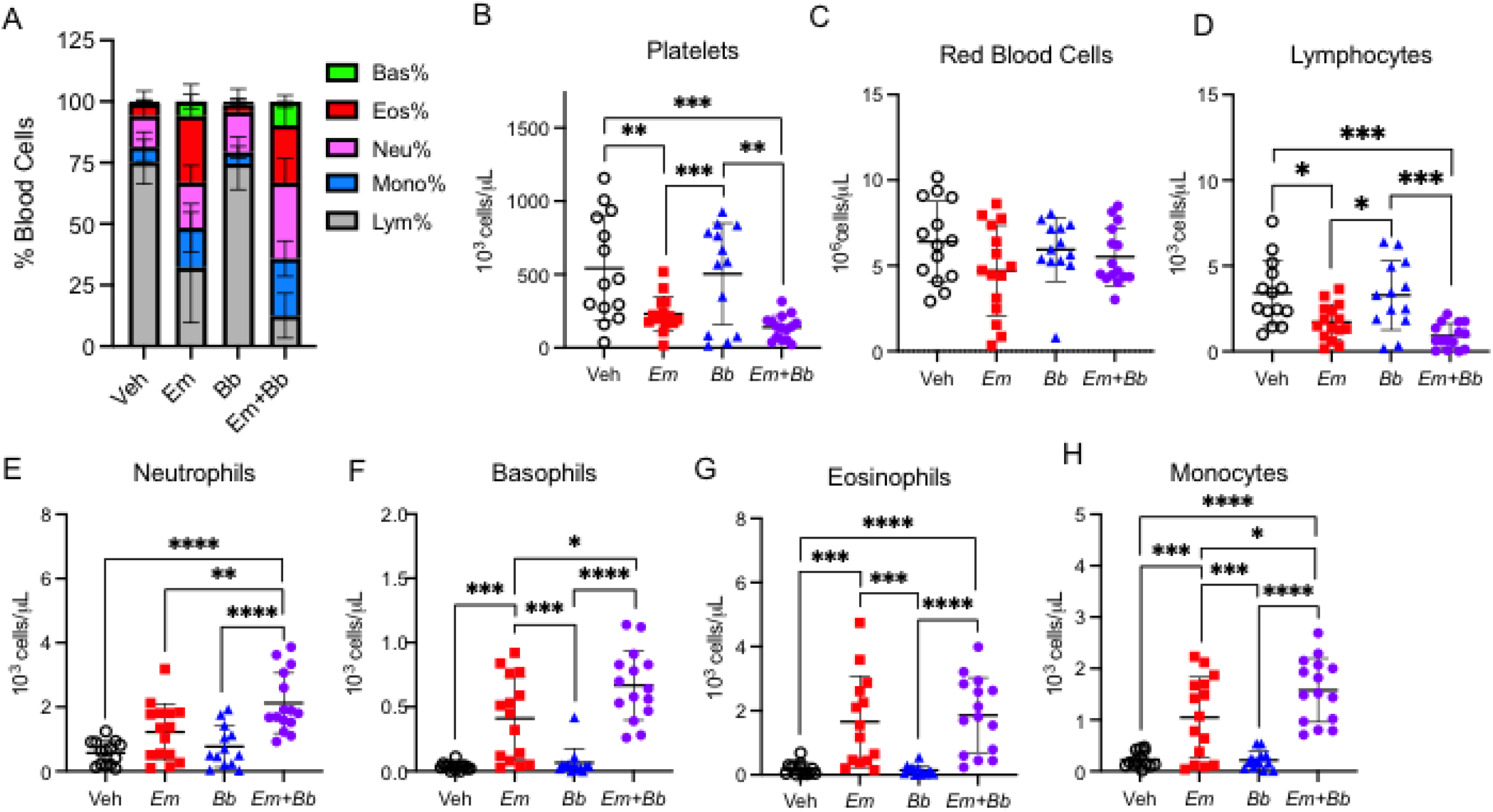
Hematological changes in co-infection with *E. muris* and *B. burgdorferi* infection. Blood was collected from infected C57BL/6Tac mice at 10 dpi. Complete blood count (CBC) analysis was performed on an automated hematology analyzer. (**A**) Composition of total white blood cells (WBC) is shown for mock-infected (SPG), *Em, Bb*, or *Em* and *Bb* co-infected mice including percent of lymphocytes, monocytes, and granulocytes (neutrophils, basophils, and eosinophils). (**B**) Platelets and (**C**) red blood cells in all groups. Total numbers of circulating (**D**) lymphocytes, (**E**) neutrophils, (**F**) basophils, (**G**) eosinophils, and (**H**) monocytes are shown. Data represent individual mice. Groups were analyzed using a one-way ANOVA with Tukey’s post-hoc comparison. Error bars represent SD. *p<0.05, **p<0.01, ***p<0.001, ****p<0.0001.

### *Em* infection drives depletion of myeloid-biased progenitor cells

To determine whether changes in the hematologic parameters of infected mice correlated with changes within the hematopoietic stem and progenitor cell (HSPC) compartment, mice were sacrificed at 10 dpi, and bone marrow was processed for flow cytometric analysis (**Supp. Fig. 4A,B**). The total pool of Lineage^−^ cKit^+^ HSPCs was reduced in the bone marrow of *Em* and co-infected mice, while Sca-1 expression increased (**Fig. 3B,C**). Phenotypically-defined HSCs (Lin^**-**^ cKit^**+**^ CD135^**-**^ CD150^**+**^ CD48^**-**^), which self-renew and give rise to more committed progenitor cells, were reduced in *Em* and co-infected animals, as were short-term-HSCs, also called multipotent progenitors (MPPs; Lin^−^ cKit^+^ CD135^−^ CD150^−^ CD48^−^; (44); **Fig. 3,D**). Among more lineage-committed HSPC populations, megakaryocyte-erythroid progenitors (MPP_Mk/E_) and myeloid-biased progenitors (MPP_GM_) (44) were depleted (**Fig. 3E,F**), whereas lymphoid-biased progenitors (MPP_Ly_) were unaltered during infection (**Fig. 3G**). Depletion of myeloid-biased MPPs correlated with the infection-induced increase in circulating myeloid cells, supporting the idea that *Em* infection induced rapid myeloid cell production, depleting progenitor cells. For all the aforementioned analyses, there were no significant differences between single-*Em* and co-infected mice, suggesting that during co-infection this phenotype is dominated by responses elicited by *Em* infection. Our data suggest that the infection-induced changes to hematopoiesis caused by *Em* may contribute to the development of Lyme arthritis in co-infected mice.

**Figure 3.**
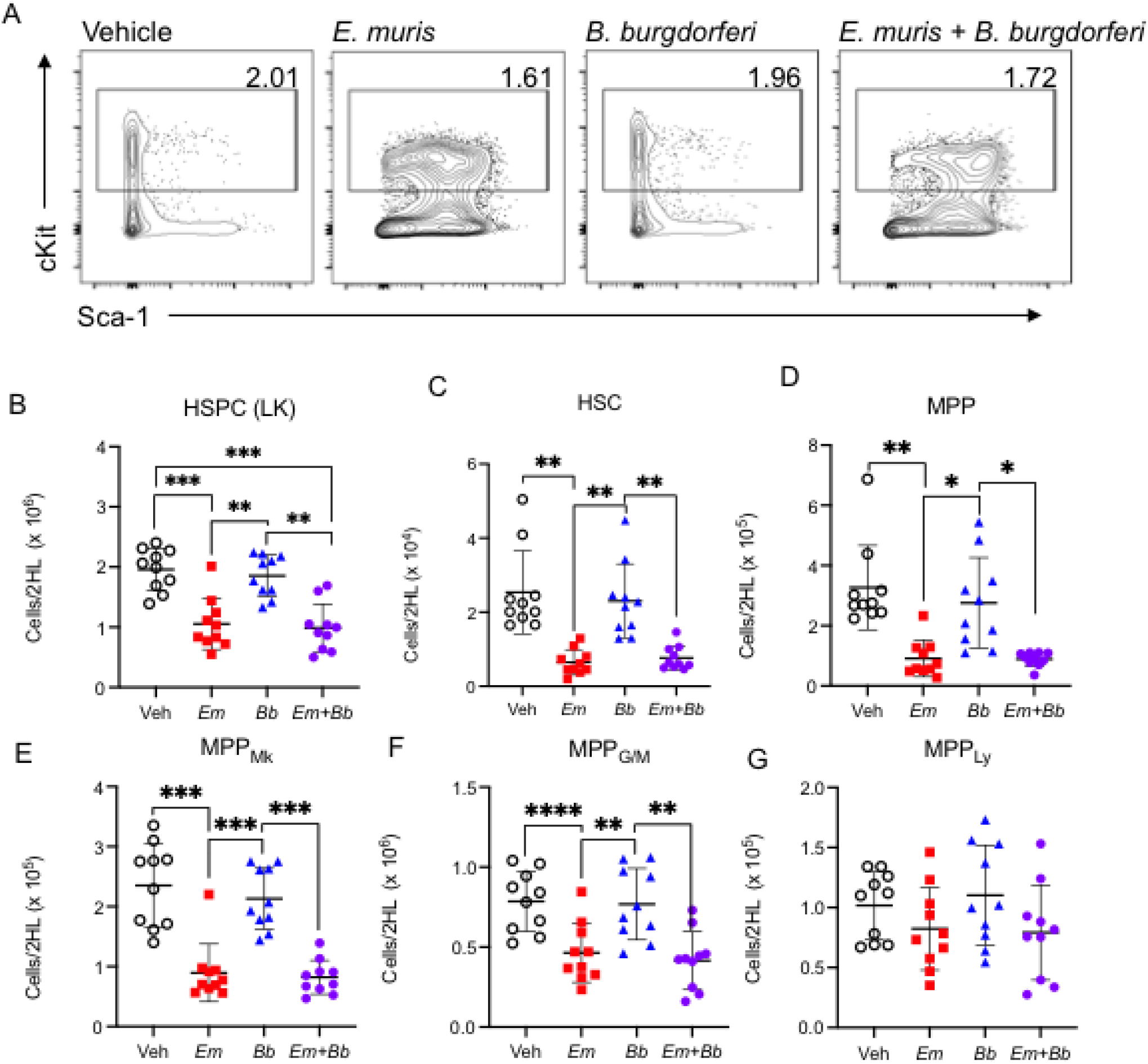
Co-infection drives loss of myeloid-biased progenitor cells in the bone marrow. Bone marrow was analyzed at 10 days post infection. (**A**) Gating shows cKit and Sca-1 expression on Lineage-negative cells and the gated region reflects the percent of cKit^+^ cells among the parent Lin^−^ gate. (**B**) Total Lin^−^ cKit^+^ HSPCs are shown for Vehicle (SPG) controls and mice infected with *Em, Bb* or co-infected both pathogens. Hematopoietic stem cells (HSC) and multipotent progenitors (MPPs) were gated based on expression of CD135, CD150, and CD48 and the absolute numbers of (**C**) HSCs, (**D**) MPPs, (**E**) MPP_Mk/E_, (**F**) MPP_GM_, and (**G**) MPP_Ly_ are shown. Data points represent individual mice. Groups were compared using a one-way ANOVA with Tukey’s post-hoc comparison. Error bars indicate standard deviation. *p<0.05, **p<0.01, ***p<0.001, ****p<0.0001.

### Co-infection results in a persistent inflammatory immune phenotype

Monocytes (CD11b^+^ Ly6G^−^ Ly6C^hi^) and neutrophils (CD11b^+^ Ly6C^−lo^ Ly6G^+^) were identified and quantified in the bone marrow by flow cytometry (**Fig. 4A**). Bone marrow cellularity of *Em*-infected mice was significantly reduced compared to control animals, consistent with previous findings (21), and bone marrow was similarly hypocellular in co-infected mice (**Fig. 4B**). Monocytes and neutrophils were depleted in the bone marrow of both *Em* and co-infected animals, whereas no change to these mature cell populations was noted in mice infected with *Bb* alone (**Fig. 4C,D**). As expected, *Em* infection induced significant splenomegaly (20, 21), which was also observed at equivalent levels in co-infection (**Fig. 4E**), and both monocytes and neutrophils were significantly increased in *Em* and co-infection (**Fig. 4F,G**). The simultaneous decrease in bone marrow myeloid cells and increase in blood and spleen myeloid cells supports the idea that *Em* infection induced rapid production and mobilization of myeloid cells, even in the context of any immune modulatory effects of *Bb* infection.

**Figure 4.**
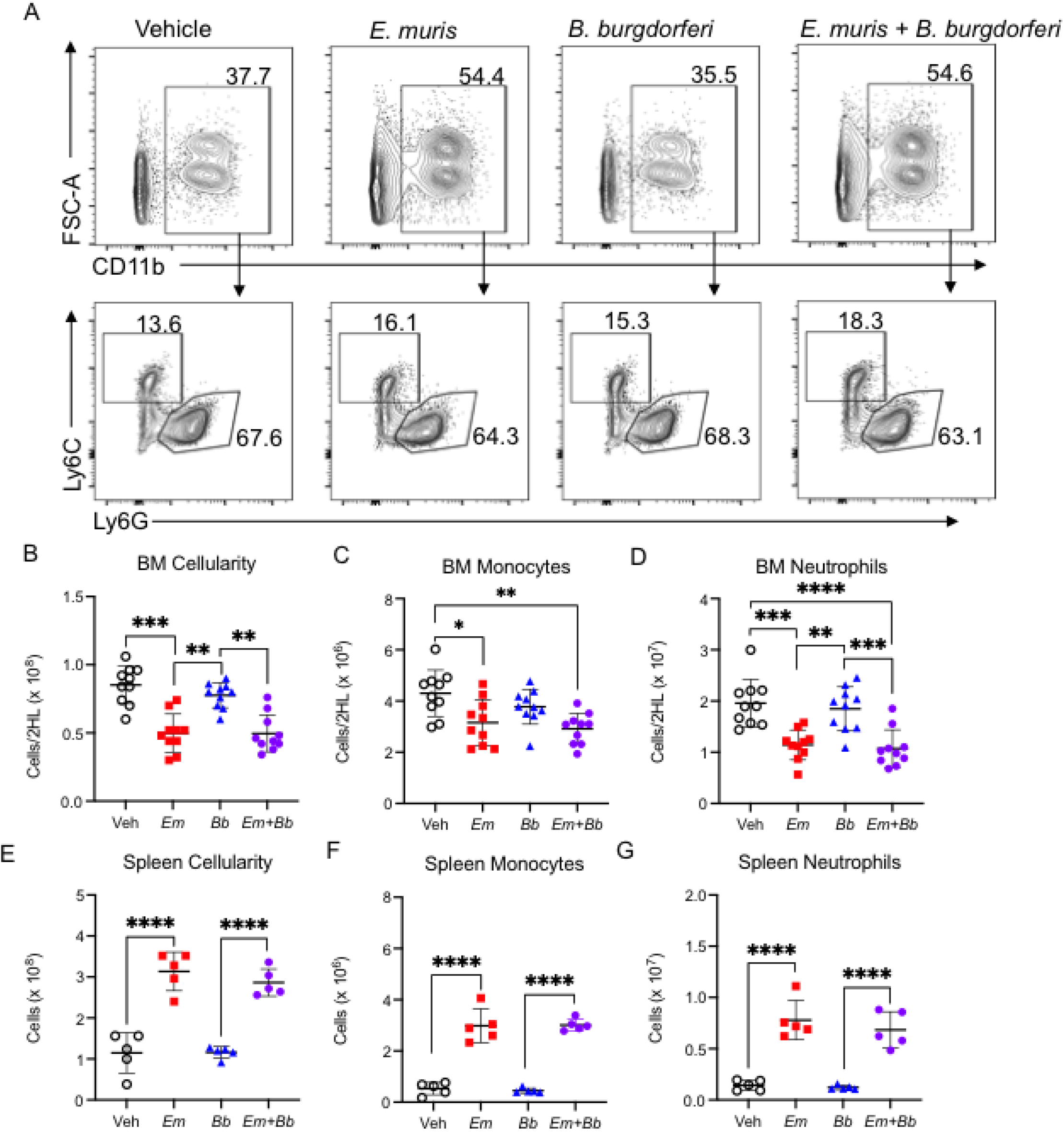
Co-infection impacts mature myeloid cells in the bone marrow and spleen at 10 dpi. Hind limbs and spleens were harvested from single- and co-infected C57BL/6Tac mice 10 days post infection. (**A**) Flow cytometric identification of CD11b^+^ cells among bone marrow cells (top row). Gating strategy to identify Ly6C^+^ monocytes and Ly6G^+^ neutrophils cells in the bone marrow of inoculated animals (bottom row). The numbers represent frequencies of gated region among CD11b^+^ cells. (**B**) Total cellularity of the bone marrow was calculated, and (**C**) monocytes and (**D**) neutrophils were quantified. (**E**) Spleen cellularity was calculated and (**F**) splenic Ly6C^+^ monocytes and (**G**) Ly6G^+^ neutrophils were quantified. Data points represent individual mice. Cell numbers were compared using a one-way ANOVA with Tukey’s post-hoc comparison. Error bars represent standard deviation. *p<0.05, **p<0.01, ***p<0.001, ****p<0.0001.

To examine systemic factors that may promote differences in hematological and immune parameters between single and co-infected animals we evaluated cytokines and chemokines in the sera. Interferon gamma (IFN-γ) was previously found to be an important driver of *Em*-induced myelopoiesis (20). We found that IFN-γ in circulation was elevated in *Em* and co-infected animals on 10 dpi but remained at baseline levels in single-*Bb*-infected mice (**Fig. 5A**), whereas TNF, IL-1β, and IL-10 were slightly elevated in the co-infected animals but not significantly different in the infected animals (**Supp. Fig. 2A-C**). The monocyte chemoattractant molecules CXCL10, CCL2, CCL4, CCL7, and CCL12 (**Fig. 5B-F**) were all elevated in the sera of *Em* and co-infected mice, consistent with the finding that ehrlichial infection results in monocytosis (45). CCL4 was significantly elevated in co-infected animals compared to singly-*Em*-infected animals while the other monocyte chemoattractants were only slightly higher in co-infected mice. Together, these data illustrate a systemic inflammatory response elicited by *Em* infection that may modulate blood cell production and migration in the context of co-infection with *Bb*.

**Figure 5.**
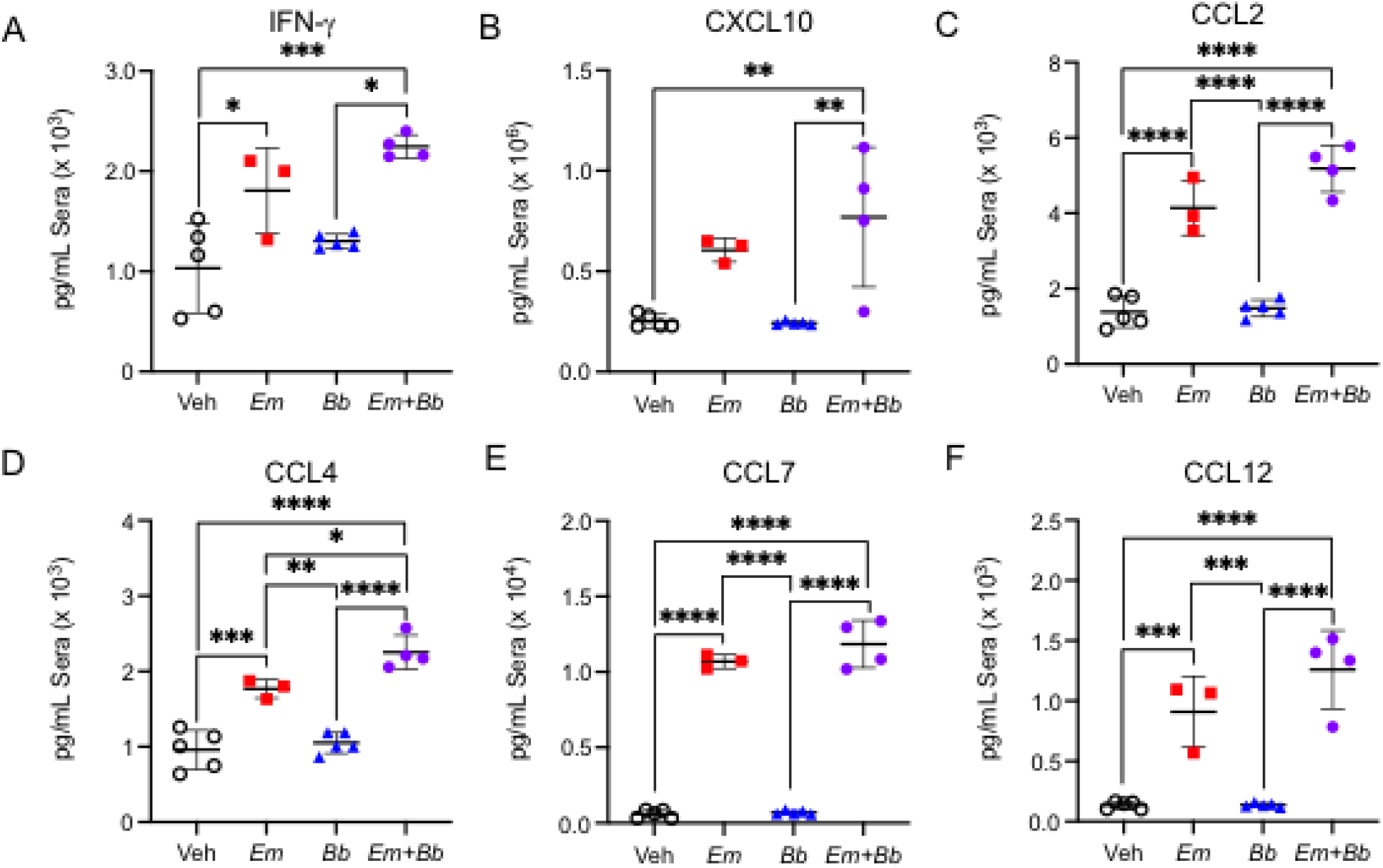
Bacterial co-infection results in an *E. muris*-dominant immune phenotype and increased monocyte chemoattractants. Immune signaling molecules were quantified at 10 dpi in the sera of single- and co-infected mice. (**A**) IFN-γ, (**B**) CXCL10, (**C**) CCL2, (**D**) CCL4, (**E**) CCL7, and (**F**) CCL12. Data points represent individual mice. Error bars indicate standard deviation. Groups were compared using a one-way ANOVA with Tukey’s post-hoc comparison. *p<0.05, **p<0.01, ***p<0.001, ****p<0.0001.

### *Em* drives a persistent increase in circulating myeloid cells

We next sought to examine the hematological parameters at day 22 post-infection, a time when mice exhibited Lyme arthritis. At this time, both co-infected and *Em-*infected mice displayed altered blood composition in comparison to healthy control mice and *Bb*-infected mice (**Fig. 6A**). Co-infected mice remained thrombocytopenic but had normal numbers of red blood cells (**Fig. 6B,C**). Neutrophils, basophils, monocytes, and eosinophils were elevated in both *Em*-infected and co-infected mice (**Fig. 6 D-G**). Unlike at 10 dpi, however, there were no significant differences among these populations between single-*Em* and co-infected mice. At 22 dpi, *Em*-induced lymphopenia was resolved in both single*-* and co-infected mice (**Fig. 6H**). Therefore, *Em* co-infection with *Bb* resulted in persistently elevated numbers of myeloid cells in the blood, correlating with the development of Lyme arthritis. Increased neutrophils were identified in the joints of co-infected mice (**Supp. Fig. 3**), further supporting the role of emergency myelopoiesis in driving development of *Bb*-induced arthritis.

**Figure 6.**
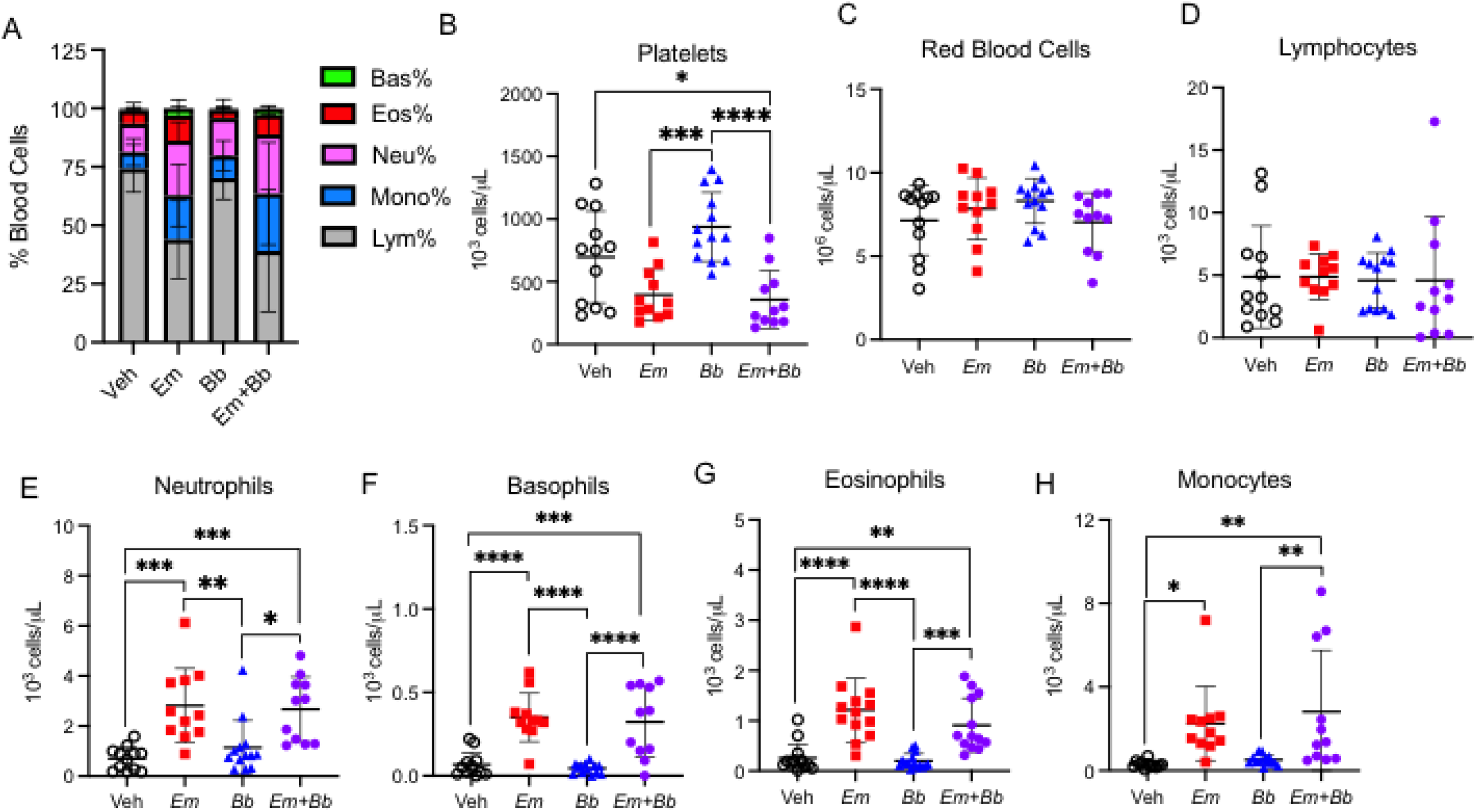
Myeloid cells remain elevated while lymphocytes return to normal in the blood. Single- and co-infected C57BL/6Tac mice were euthanized at 22 dpi, when arthritis is prominent, and CBC analysis was performed on an automated hematology analyzer. (**A**) Composition of total white blood cells (WBC) is shown for mock-infected (SPG), *Em, Bb*, or *Em* and *Bb* co-infected mice including percent of lymphocytes, monocytes, and granulocytes (neutrophils, basophils, and eosinophils). (**B**) Platelets and (**C**) red blood cells in all groups. Total numbers of circulating (**D**) lymphocytes, (**E**) neutrophils, (**F**) basophils, (**G**) eosinophils, and (**H**) monocytes are shown. Data represent individual mice. Groups were analyzed using a one-way ANOVA with Tukey’s post-hoc comparison. Error bars represent SD. *p<0.05, **p<0.01, ***p<0.001, ****p<0.0001.

### Co-infection drives persistent inflammation in the bone marrow

To evaluate the inflammatory state of the hematopoietic compartment when Lyme arthritis was present, we examined the bone marrow at 22 dpi. IFN-γ levels were significantly increased in the bone marrow of co-infected mice (**Fig. 7A**), as were the chemokines CXCL10 and monocyte chemoattractants CCL2, CCL4, CCL7, and CCL12 (**Fig. 7B-F**). Singly-*Em*-infected mice did not have elevated IFN-γ at day 22, however their bone marrow was characterized by elevated chemokines, including CXCL10, CCL12, CCL4, and CCL5. Notably, co-infected mice had significantly elevated CCL7 and CCL12, relative to mice singly infected with *Em*. Thus, whereas *Em* infection alone was characterized by moderate and persistent changes to peripheral blood that were similar to co-infected animals, the bone marrow compartment exhibited a unique inflammatory signature as compared *Em*-infected mice. These data suggest a correlation between the co-infection arthritis phenotype and IFN-γ, CXCL10, CCL2, CCL7, CCL12, and CCL4 responses.

**Figure 7.**
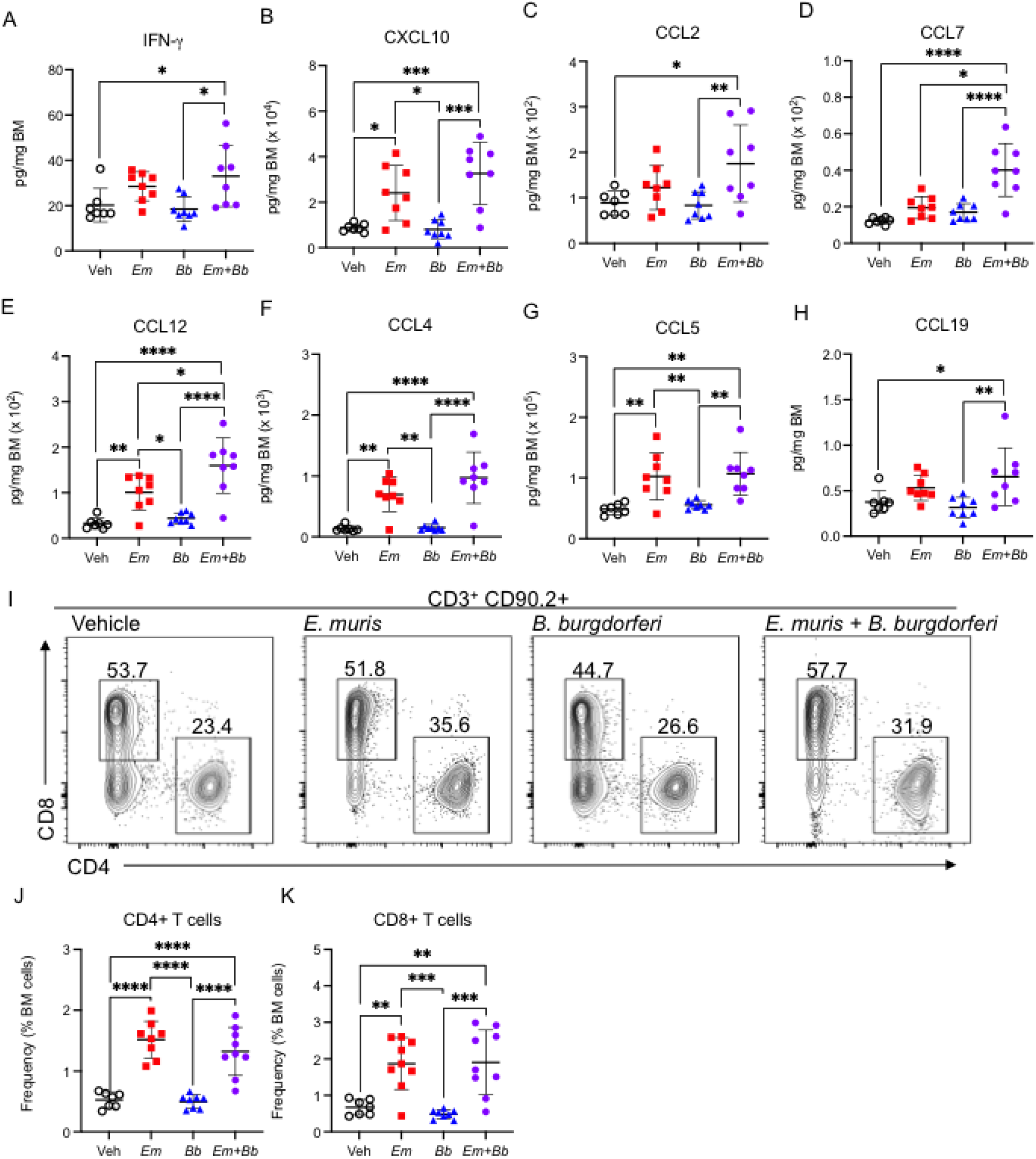
Co-infection drives persistent inflammation in the bone marrow. Co-infected mice were euthanized, and bone marrow processed for protein and cell analysis via flow cytometry. Inflammatory cytokines were analyzed and concentrations of bone marrow cytokines and chemokines are shown including (**A**) IFN-γ, (**B**) CXCL10, (**C**) CCL2, (**D**) CCL4, (**E**) CCL7, and (**F**) CCL12, (**G**) CCL5, and (**H**) CCL19. Concentrations were normalized to total bone marrow protein. (**I**) Flow cytometric gating of CD90.2^+^ CD3^+^ lymphocytes for CD4+ or CD8+ T cells. The number above each gated region is the percent of parent gate for representative mice in each group. (**J**) Absolute frequencies of CD90.2^+^ CD3^+^ CD4^+^ T cell and (**K**) CD90.2^+^ CD3^+^ CD8^+^ T cells among total bone marrow cells is shown. Data points represent individual mice. Groups were compared using a one-way ANOVA with Tukey’s post-hoc comparison. Error bars represent standard deviation. *p<0.05.

Furthermore, CCL5 and CCL19 were also elevated in co-infected mice at 22 dpi (**Fig. 7G,H**), with CCL19 elevated only in the context of co-infection. As CCL19 is a chemotactic signal for T cells we evaluated the lymphocyte compartment in the bone marrow. CD3^+^ CD90.2^+^ cells were gated on CD4 and CD8 expression to identify CD4+ and CD8+ T cells (**Fig. 7I**). Significantly elevated frequencies of both CD4 and CD8 T cells were observed in the bone marrow of both *Em* and co-infected mice (**Fig. 7J,K**), suggesting the *Em*-driven inflammatory phenotype seen in mice at 10 dpi persists to 22 dpi and promotes accumulation of lymphocytes in the bone marrow. Together our data suggest that *Em* infection sensitizes mice to a more pronounced inflammatory state upon co-infection with *Bb*, supporting the conclusion that persistent inflammation in the bone marrow may play a key role of in driving the development of Lyme arthritis.

## Discussion

Tick-borne diseases are increasing in prevalence throughout the United States, posing a significant public health challenge. Lyme disease, caused by *B. burgdorferi* (*Bb*), is the most common tick-borne disease in the US (3, 46). In this study we show that co-infection of B6 mice with both *Bb* and an *Em* results the development of severe arthritis. BALB/c and C57BL/6 are resistant to the development of Lyme arthritis while C3H mice develop arthritis and carditis consistent with human patients (39). Crandell et al. showed that C57BL/6 mice have reduced expression of IFN-induced genes when compared to C3H mice, and elevated expression of genes associated with tissue repair (47). Herein, infection with *Em* resulted in transient activation of HSCs, which has previously been shown to be IFN-γ-dependent process and associated with production of monocytes and granulocytes (20). IFN-γ is critical for control of *Em* (21, 48), thus, these responses are likely an adaptation of the host to control *Em* infection. IFN-γ production occurs during *Bb* infection in humans, with the cytokine identified in the blood and cerebrospinal fluid of infected individuals, and high levels of IFN-γ are associated with long-lasting inflammation and antibiotic-refractory arthritis (49, 50). Consistent with Crandall et al., (47) we show that B6 mice infected with *Bb* alone did not produce IFN-γ, and did not develop Lyme arthritis. However, during coinfection of *Bb* with *Em*, B6 mice produced high levels of IFN-γ and experienced profound infiltration of neutrophils and lymphocytes into the ankle joint. Together, these data suggest that the presence of *Em*-elicited IFN-γ during *Bb* infection of B6 mice drives the development of Lyme arthritis in otherwise resistant mice.

*Em* causes significant hematologic disturbances (14), and the hematologic disturbances observed in *Em* and *Bb* co-infected mice mirrored single *Em*-infected animals, with profound monocytosis at 10 dpi, the peak of *Em* infection (19). This *Em*-dominant immune response persisted at 22 dpi, the relative peak of *Bb* infection (8). At 22 dpi, lymphopenia resolved in all infected animals, while granulocytosis remained prominent. Supporting the idea that granulocytes contribute to arthritis is the observation of neutrophils in the joint lesions. C57BL/6 mice infected with *Bb* exhibited no detectable disturbances to their hematologic parameters, and resembled vehicle-treated mice, suggesting that the organism evades or suppresses host immune responses. Co-infected mice mirrored *Em* infection with respect to inflammation and myelopoiesis and correlated with severe Lyme arthritis. This prompted us to question whether *Bb* growth *in vivo* was enhanced by *Em* co-infection which we reasoned could support persistence of *Bb*. This raised the possibility that *Bb* infection alone failed to induce arthritis because it was unable to establish infection. To rigorously test whether single infection with *Bb* was able to establish infection in B6 mice, and to determine the impact of co-infection on *Bb* persistence, we performed a xenodiagnosis assay, a robust method to determine spirochete infection in mammals (40, 51). We found that mice infected with *Bb* alone or co-infected with both *Bb* with *Em* transmitted *Bb* to larval ticks demonstrating that all mice were indeed infected with *Bb*. These data confirm that the failure of *Bb* infection alone to induce development of arthritis was not due a failure of the pathogen to persist in B6 mice and demonstrate the role of *Em* in driving an inflammatory response that promotes arthritis. These data demonstrate that despite the ability of single-*Bb* to persist in C57BL/6 mice and transmit to naïve ticks, these mice exhibited no joint pathology or hematologic disturbances. It is a well-documented phenomenon that *Bb* evades or suppresses host immune responses during infection. For example, expression of CspA, CspZ, and OspE-related proteins act to inhibit the alternative pathway of complement activation to prevent cell lysis and inflammation (10). Thus, it is unlikely that *Bb* itself is sufficient for joint inflammation, but rather, host factors play a key role in determining disease progression and such factors can be altered by the co-infection of other tickborne pathogens. The hematologic parameters observed in murine co-infection were consistent with those seen in human co-infection with *Bb* and Ehrlichia species (17) further supporting the clinical relevance of our findings. Determining the mechanisms that result in recruitment to the joint tissue in co-infected animals will be critical for treating joint pathology in Lyme disease patients.

*Bb* and ehrlichial pathogens are transmitted to a host via tick feeding. Tick bites transmit not only bacteria but also salivary proteins (salivary gland extracts, or SGE). In the context of *Bb* transmission, tick salivary proteins aid in establishment of infection through suppression of local host immune mechanisms (52). The *Ixodes scapularis* protein Salp15 has been shown to protect *B. burgdorferi* from antibody-mediated killing in a murine model of Lyme disease (53). Additionally, *Ixodes*-produced Salp14 serves as an anticoagulant that promotes the length of tick feeding (54). Tick feeding delivers the pathogen into the dermis (intra-dermal), with initial immune responses to the pathogen occurring within this tissue. Sertour et al. found that significant changes in tissue tropism occurred between mice that were infected via needle inoculation and tick feeding (38), therefore it stands to reason that the i.p. route of pathogen administration used here may play a role in the resulting immune response. However, regardless of pathogen presence, tick feeding results in a localized host immune response, likely introducing additional variables that may impact disease outcomes (54). Therefore, future work is warranted to evaluate specific impacts of tick feeding on bacterial persistence and arthritis development in the context of co-infection.

When *Em* infection occurs simultaneously with *Bb* mice normally resistant to Lyme arthritis develop severe joint pathology by 3 weeks. Our data are consistent with previous findings that co-infection of *Bb* with other tick-borne pathogens exacerbates the severity of Lyme disease (15, 55, 56). A myriad of immune cells and cytokines were altered in the blood and bone marrow of co-infected mice that were unaltered in single *Bb*-infected mice. The observation that co-infection elicited more pronounced inflammatory changes during early stages of co-infection (10 dpi) suggested an interaction between the host and co-infecting pathogens that permitted development of arthritis. The tropism of the spirochete for joints and the increased presence of inflammatory cells in the blood of coinfected mice enabled neutrophils and monocytes to home to the joint, driving arthritis. Further studies are needed to identify which cell types and signaling molecules are necessary to initiate and perpetuate the disease phenotype.

## Acknowledgements

The authors would like to thank Hui Jin Jo and Allison Seyfried for technical assistance.

## Author contributions

JLB and ALM performed experiments, analyzed data, and wrote the paper. SRT designed and performed experiments and analyzed data. JRD performed experiments and analyzed data. XY and UP performed histopathology analysis. TN Performed experiments. YPL designed experiments and analyzed data. KCM conceived the project, designed experiments, analyzed data, and wrote the paper.

